# Social isolation is associated with microRNA expression changes in the visual system of poison frog tadpoles

**DOI:** 10.64898/2026.09.04.749321

**Authors:** Neil M. Khosla, Max M. Madrzyk, Keira Nakamura, Mei Li Gallardo Palmeri, Julie M. Butler, Lauren A. O’Connell

## Abstract

Brain microRNA (miRNA) expression can shift in response to the social environment, translating external cues into regulatory action that can influence brain structure, neural activity, and behavior. miRNAs are key drivers of behavioral flexibility, regulating gene networks involved in neuroplasticity. We manipulated social rearing conditions of brilliant-thighed poison frog tadpoles (*Allobates femoralis*) to test whether isolation-linked behavioral changes are associated with altered miRNA expression in the midbrain and eyes, tissues linked by the retinotectal tract, a known connection for trafficked precursor miRNAs. We asked whether isolation alters visually guided behavior using a light/dark preference assay and found that isolated tadpoles spent less of the trial on the dark side than did group-housed tadpoles. In a social place preference assay, group-housed tadpoles preferred a zone near a conspecific over an object control when provided with multimodal sensory access, but not when access was restricted to vision alone. We next tested whether social rearing conditions influenced miRNA expression and found that isolated tadpoles exhibited upregulation of miR-15-P1c_3p*, the canonically non-dominant arm of an understudied miRNA, in the midbrain and eyes. We detected a shift in arm dosing for miR-124, a miRNA with conserved roles in neural differentiation and plasticity. Finally, social isolation during development was associated with downregulation of parathyroid hormone 2 (*pth2*), a peptide associated with social isolation and mechanosensation in other aquatic larvae. This work contributes to a growing body of literature implicating miRNAs in developmental neuroplasticity, presenting precedented and novel signatures of the transcriptional and regulatory response to isolation.

## Introduction

In group-living species, the social environment provides a range of benefits during development. Some are passive, like safety in numbers (Foster, 2017), while others are active, providing a setting for learning cooperative and competitive skills (Pellis et al., 2023). Isolation can therefore be a profoundly disruptive stimulus for a social animal, leading to wide-ranging effects such as increased aggression in rodents (Takahashi, 2025), altered social sensitivity in zebrafish (Anneser et al., 2020), and disruption of immune function in social insects (Meunier, 2015). Studying developmental isolation in social species can reveal how social and sensory processes are driven by neural systems at multiple levels, from broad brain regions to specific molecular mechanisms enabling flexible responses to social stimuli. Though the impact of social experience is often studied in the adult brain, the influence of isolation on regulatory mechanisms in the developing brain is relatively understudied.

MicroRNAs (miRNAs) are molecular factors that translate social experiences into coordinated changes in gene expression. These short, non-coding RNAs post-transcriptionally regulate gene expression by guiding the RNA-induced silencing complex (RISC) to binding sites in 3’UTRs of target transcripts (Bartel, 2009). Brain miRNA expression changes in response to the social environment, impacting gene expression profiles and behavior. For example, isolation-induced aggression in mice is related to regulation of brain-derived neurotrophic factor (*bdnf*) by miR-206 in the ventral hippocampus (Chang et al., 2020). Kin recognition in *Xenopus* tadpoles is linked to miRNA-mediated regulation of neurotransmitter switching in the accessory olfactory bulb (Dulcis et al., 2017). Since miRNAs regulate networks of genes, slight shifts in the expression of a single miRNA can have broad-reaching effects. For instance, in locusts, social phase polyphenism is linked to regulation of multiple genes in a dopamine biosynthesis pathway by miR-133 (Yang et al., 2014). These studies reveal shared function for miRNAs in behavior across diverse taxa, forming the basis for an emerging, interdisciplinary field.

One underexplored layer of this regulation is arm selection. Each miRNA hairpin contains two alternative mature sequences, the 5p and 3p arms. After Dicer releases the mature miRNA duplex from the hairpin, one arm is preferentially loaded into RISC in a process called arm selection. Since the two arms carry distinct seed sequences, arm selection determines which targets a given miRNA regulates (Pinhal et al., 2025). In multiple types of cancer, alternate arms of the same miRNA are inversely expressed and target transcripts that play opposing roles in progression of the disease (Ren et al., 2018; Zhang et al., 2019). However, this phenomenon has yet to be identified in the nervous system in response to social experience.

Poison frog species (family Dendrobatidae) exhibit diversity in species-specific social strategies, ranging from cannibalistic to gregarious (O’Connell, 2020). Tadpoles are transported on the backs of parent frogs to pools of water, which vary in size, nutrient availability, and conspecific density across species. For example, poison frogs of genus *Ranitomeya* transport their young singly to resource-poor pools, where tadpoles display aggressive strategies (Brown et al., 2009). Other species exhibit flexibility in social settings, like the brilliant-thighed poison frog, *Allobates femoralis* (Boulenger, 1884), which transports tadpoles to nutrient-rich pools where tadpoles usually develop in groups (Ringler et al., 2013). Although visual systems in tadpoles are immature compared to adults (Butler et al., 2024), visually guided behavior is already shaped by the rearing environment (Fouilloux et al., 2023). In *A. femoralis*, conspecific visual cues alone are sufficient to increase tadpole activity (Szabo et al., 2021). The neuroethological context for *A. femoralis* social rearing plasticity makes it a fitting model for investigating context-dependent regulation of gene expression.

Successful behavioral adaptation relies on coherent communication between sensory organs and the brain. The eye-brain axis is a well-described substrate for molecular regulation of visually motivated behaviors and the role of miRNAs in this process is beginning to be uncovered in the developing brain. In *Xenopus* tadpoles, miR-124 (a conserved regulator of neural differentiation and plasticity (Olde Loohuis et al., 2012)) and miR-181 are important for accurate axon targeting during early visual development (Baudet et al., 2012; Corradi et al., 2020). That work established the visual system as a model for small RNA regulation of neuronal wiring and function. Whether the environmental sensitivity of brain miRNA expression extends to this axis is unknown. Here, we tested the hypothesis that social isolation of *A. femoralis*, a normally group-reared species, would alter visually guided behavior and expression of miRNAs and mRNAs in eyes and brain regions involved in visual processing.

## Methods

### Experimental animals

*Allobates femoralis* adult frogs were purchased from Indoor Ecosystems (Whitehouse, Ohio). All *Allobates femoralis* tadpoles used in this study were bred in our captive poison frog colony at Stanford University. Briefly, 3-5 males and females were housed together in glass terraria containing moss substrate, live plants, multiple shelters, egg deposition sites (petri dishes), and multiple water pools for tadpole deposition. Frogs were housed on a 12 h:12 h light:dark cycle and fed fruit flies dusted with vitamin supplements. Water pools were checked regularly for transported tadpoles, which were subsequently moved to circular containers outside of the tank.

Tadpole clutch size ranged from 12-16 individuals, and each clutch was split so that half of the clutch was reared as a group (n ≥ 5 per group), and the rest were reared in isolation. Tadpoles were fed an excess of brine shrimp flakes and tadpole pellets (Josh’s Frogs, Owosso, MI, USA) three times per week. Cup water was refreshed twice per week. Each rearing container contained sphagnum moss and tea leaves as nutrient supplements. Tadpole morphology (body mass, body length, body width) was measured immediately before dissections. All tadpoles were raised until Gosner stages 28-32 (early hindlimb development) (Gosner, 1960), and paired animals were sampled the same day, at which point they were used for the various experiments described below. All procedures were approved by the Stanford Administrative Panel on Laboratory Animal Care (protocol no. 33097).

### Light/dark phototaxis assay

To assess tadpole visual capabilities and light-environment preferences in tadpoles, we performed a phototaxis assay designed in our lab (Butler and O’Connell, 2023). Briefly, the behavioral arena consisted of a petri dish (10 cm) seated on the lid of a larger petri dish (15 cm), whose bottom was half-painted opaque black, creating adjacent light and dark halves beneath the tadpole. The painted dish was mounted such that it could be rotated to flip the light/dark sides without disturbing the tadpole.

We tested stage matched (Gosner 28-32) tadpoles reared either in social isolation or group housing (n = 8 group-housed, n = 8 socially isolated). Ten minutes before trials, tadpole cups were moved to the procedure room to acclimate. The arena was filled with 40 ml of water and focal tadpoles were placed in the center. Behavior was recorded from above with a GoPro. Tadpoles explored the arena for 3 minutes, at which point the light/dark sides were flipped, and behavior was recorded for an additional 3 minutes. Each video was manually scored for time spent on each side (light/dark).

### Social place preference

Social place preference was assayed in two separate experiments differing in stimulus chamber barrier type: perforated barriers that permitted water exchange between chambers or solid barriers that permitted visual access only. Different tadpoles were used for each barrier type, with one tadpole from each clutch per housing treatment in each experiment (n = 8 clutches). Focal animals were stage matched at Gosner 28-32. Trials were run in U-shaped social place preference arenas (Nakamura et al., 2024) containing 50 ml of water. Each trial began with a 30-minute acclimation with both stimulus chambers empty, which also provided a within-trial baseline to test innate side bias (tadpoles did not exhibit preference at baseline; Fig. S1). Stimuli were then introduced: an unrelated conspecific tadpole on one side and a hexagonal nut (object control) on the other, with side randomized across trials. Behavior was recorded for 60 minutes following stimulus introduction.

Tadpoles were tracked using DeepLabCut, a tool used for automated behavioral tracking (Lauer et al., 2022; Mathis et al., 2018). A ResNet152 multi-animal model was trained on 1600 manually labeled frames. Region of interest (ROI) boundaries were defined as x-y coordinates on video stills in ImageJ, and a point-in-polygon test assigned the focal tadpole to a zone in each frame based on its proximity to the stimulus chambers.

### Small RNA and mRNA library preparation

Tadpoles were euthanized using an overdose of benzocaine, 24 hours after social place preference experiments, and used to prepare RNA libraries. Immediately after, brains were dissected (forebrain, midbrain, and hindbrain) and these samples, along with the eyes, were placed into separate tubes (Z763799, Millipore Sigma) and snap-frozen on dry ice. Tissues were pooled (n = 3 individuals per clutch, per condition) to ensure sufficient tissue for minimum RNA input requirements in library preparation protocols.

All tissue samples were stored at −80°C until total RNA was extracted from both midbrain and eye tissues using the Monarch Total RNA Miniprep Kit (T2010, New England Biolabs) following the manufacturer’s protocol for low-input extraction with mechanical perturbation. Samples were assigned to extraction batches at random with respect to both housing condition and clutch. High-quality RNA samples from midbrain and eye (RIN > 7) were used for library preparation.

For small RNA libraries, 24 ng of input RNA were used with the NEBNext Multiplex Small RNA Library Prep Kit (E7330L, New England Biolabs) according to the kit instructions. The prepared libraries underwent a two-step purification process. Initial purification was performed using the Monarch PCR & DNA Cleanup Kit (5 μg) (T1030S, New England Biolabs), followed by further size selection to isolate microRNA-sized libraries using the PippinHT system (HTG3010, Sage Science). The purified and size-selected libraries underwent QC with Qubit and Agilent TapeStation before equimolar pooling of libraries and subsequent final QC of pooled libraries using Qubit HS DNA and Agilent BioAnalyzer. Pooled libraries were sequenced by Azenta Life Sciences (GENEWIZ) on an Illumina NovaSeq X in 2×150 bp paired-end mode. Libraries were sequenced in two runs (the second run to increase read depth) and reads from both runs were concatenated per sample prior to analysis. Since small RNA inserts are shorter than the read length, only Read 1 (the forward read) was used for downstream alignment.

mRNA libraries were prepared using 50 ng of input RNA in the NEBNext Ultra II Directional Library Prep Kit for Illumina (E7760L). The NEBNext Poly(A) mRNA Magnetic Isolation Module (E7490) was used to isolate polyadenylated RNAs. The protocol proceeded following manufacturer instructions. Libraries were quantified using the NEBNext qPCR Library Quant Kit for Illumina (E7630S). mRNA libraries were sequenced by Azenta Life Sciences (GENEWIZ) on an Illumina NovaSeq X in 2×150 bp paired-end mode.

### Sequence alignment and expression quantification

Small RNA reads were processed with sRNAbench (sRNAtoolbox) (Aparicio-Puerta et al., 2022), mapping reads against a *Xenopus tropicalis* mature miRNA reference. Reference entries with identical mature sequences were collapsed to a single entry and are reported under the first member of each group. miRNA names follow MirGeneDB nomenclature (Clarke et al., 2025) with the *Xtr*-species prefix omitted; an asterisk denotes the non-dominant (“star”) arm. We created a miRNA annotation against the *A. femoralis* genome using MirMachine (Umu et al., 2023), but it is computationally predicted, unvalidated, and does not discern paralogs. We therefore mapped to the most closely related manually curated miRNA reference (*X. tropicalis*), restricting detection to conserved, mature miRNAs. Adapters were trimmed with cutadapt v1.18 (adapter: AGATCGGAAGAGCACACGTCTGAACTCCAGTCAC) with a quality cutoff of Q20. Reads were aligned with bowtie, allowing up to one mismatch across the full read (alignType=v, noMM=1). Given the deep divergence between *A. femoralis* and *X. tropicalis* (∼190 Myr (Hime et al., 2021)) this restricts detection to strongly conserved miRNAs. Counts were summed across isomiRs to the mature miRNA level and reads mapping to multiple references contributed 1/n to each of the n references matched. mRNA library reads were trimmed for quality and adapters with cutadapt v1.18 (Q20, minimum length 36 nt), and aligned to the *A. femoralis* genome [ASM3357653v1, GCA_033576535.1] with STAR v2.7.10b (Dobin et al., 2013). Gene-level counts were generated with STAR, taken from the reverse-stranded column. Multimapping reads were excluded from gene-level counts.

For each dataset, counts were assembled into per-library matrices. Differential expression was performed on raw counts in DESeq2 (Love et al., 2014) and model designs are provided in *Statistical analyses*. For the heatmap, counts were variance-stabilized (DESeq2) and z-scored within each miRNA.

### miR-124 precursor annotation and structure

miRNA loci were predicted *de novo* in the *A. femoralis* genome assembly with MirMachine v.0.2.13 (Umu et al., 2023) using the *Xenopus* node covariance models, recovering three miR-124 loci. Mature 5p and 3p sequences are identical across miR-124 paralogs, which differ only by 1-2 nucleotides in the terminal loop, so reads cannot be attributed to individual paralogs. Thus, counts of miR-124 arms are reported as family totals. The precursor used for structure prediction (miR-124-P1-v1) was the one with the most reads spanning its terminal loop. Minimum-free-energy secondary structure was computed with RNAfold (ViennaRNA (Lorenz et al., 2011)).

### Statistical analyses

All statistical analyses were performed in R (v4.4.1).

In all box plots, boxes show the median and interquartile range. Whiskers extend to the most extreme value within 1.5 × the interquartile range.

*Morphology*: Body length, width, and weight were compared between groups and paired by clutch (n = 6 clutches). For each measure, we used a paired t-test on the per-clutch differences (isolated – grouped).

### Phototaxis preference

For the phototaxis assay, the response was the proportion of the full trial spent on the dark side. Since group-housed siblings share a rearing cup, group-housed values were averaged to cup mean per clutch, whereas socially isolated tadpoles are singly housed and contributed individual values (n = 4 group-housed cups with 2 tadpoles assayed from each; n = 8 isolated tadpoles). Housing treatments were compared with a two-sided Welch’s t-test. As a secondary analysis, results from a two-sided Wilcoxon rank-sum test are reported for comparison (W = 28, p = 0.050). Each condition was compared against chance (0.5) with a two-sided one-sample t-test.

### Social Place Preference

Point-in-polygon tests assigned the focal tadpole to a social zone, an object zone, or the connecting hallways on each tracked frame. Time in hallway was excluded from analysis. Analyses only include clutches with all four housing × barrier-type conditions present (n = 8 clutches, 32 tadpoles per phase). Time in the two stimulus zones was compared within tadpole, separately for each barrier type, housing condition, and phase. Occupancy was expressed as minutes in each zone within the 60-minute trial window and compared with a linear mixed model (time ∼ zone) with a random intercept for animal, fitted by maximum likelihood. χ^2^ values are likelihood-ratio tests of the full model against null. P-values were Benjamini-Hochberg corrected across barrier type (perforated/solid) × rearing (grouped/isolated) × phase (baseline/trial) comparisons.

### Differential expression analyses

Features were retained for testing if their mean count across libraries was at least 5, or their count exceeded 11 in at least 20% of libraries. mRNA and miRNA differential expression were tested with DESeq2 with the Wald test on raw counts, contrasting tissues (eyes vs. midbrain) or housing condition libraries within each tissue. Size factors were estimated with the “poscounts” method. Models were fit as ∼clutch + tissue and ∼clutch + housing. RNA extraction batch was randomized with respect to housing and clutch, so it was not included as a covariate. Dispersion was estimated with the default “parametric” fitType for mRNA, and the “local” fitType for miRNA libraries, due to the relatively smaller number of miRNA features tested. Log_2_ fold changes were moderated with ashr (Stephens, 2016), and fold change thresholds were applied to the moderated estimates. Features with p_adj_ < 0.05 (Benjamini-Hochberg) and |log_2_ fold change| > 1.5 (mRNA) or > 0.5 (miRNA) were considered differentially expressed. A lower threshold was applied to miRNAs since they act on many targets, and we therefore treat the miRNA threshold as an exploratory screening criterion rather than an effect-size claim.

### Arm dosage

For each tissue, miRNA hairpins with both arms expressed (each arm ≥ 5 reads summed across libraries and detected in ≥ 20% of libraries) were tested for housing-dependent arm expression. Arm-dosage analyses used the same paired-clutch cohort as the housing contrasts. For each hairpin, the per-library log_2_ 3p/5p ratio was modeled with a linear model (log_2_ ratio ∼ clutch + housing). P-values were Benjamini-Hochberg corrected for multiple hypothesis testing, and hairpins with p_adj_ < 0.05 considered to show housing-dependent arm dosing. As a complementary test, per-library ratios were averaged within clutch and compared between housing conditions with a clutch-paired Wilcoxon signed-rank test. For significant hairpins, effect sizes (Hedges’ g and Cliff’s δ) were calculated on arm ratios.

### pth2 3’UTR recovery

Since *pth2* had no annotated 3’UTR, we recovered one from our own data. All midbrain and eye libraries were pooled and the 3’ end of the transcript was identified from reads carrying poly(A) tails not present in the genome (mean coverage 519× across the 3’UTR). The recovered 81 nt of sequence was searched for miRNA binding sites (6mer, 7mer, and 8mer seed matches). It is not uncommon for 3’UTRs of small peptides to be short, and the 3’UTR for the human ortholog of *pth2* is comparably small at 49 nt. The 15 nt annotated for the ortholog in the golden poison frog, *Phyllobates terribilis* (Myers, Daly, and Malkin, 1978), is near-identical to the start of the recovered *A. femoralis* sequence.

## Results

## Social isolation reduces dark side preference in a phototaxis assay

We first tested whether social rearing conditions influence responses to non-social visual cues by comparing group-housed and socially isolated tadpoles in a phototaxis light/dark assay (Fig. 1a-b), which is used to assess exploratory behavior and visual sensitivity (Adebogun et al., 2023; Maximino et al., 2010). We found that group-housed and socially isolated tadpoles differed in visually guided behavior. Isolated tadpoles spent a smaller proportion of the trial on the dark side than did group-housed tadpoles (Fig. 1c; Welch’s t-test, t_9.5_ = 2.48, p = 0.034). Relative to chance, group-housed tadpoles spent more of the trial on the dark side (one-sample *t*-test vs. 0.5, t_3_ = 4.50, p = 0.020), whereas isolated tadpoles did not differ from chance (t_7_ = 0.50, p = 0.631).

**Figure 1.**
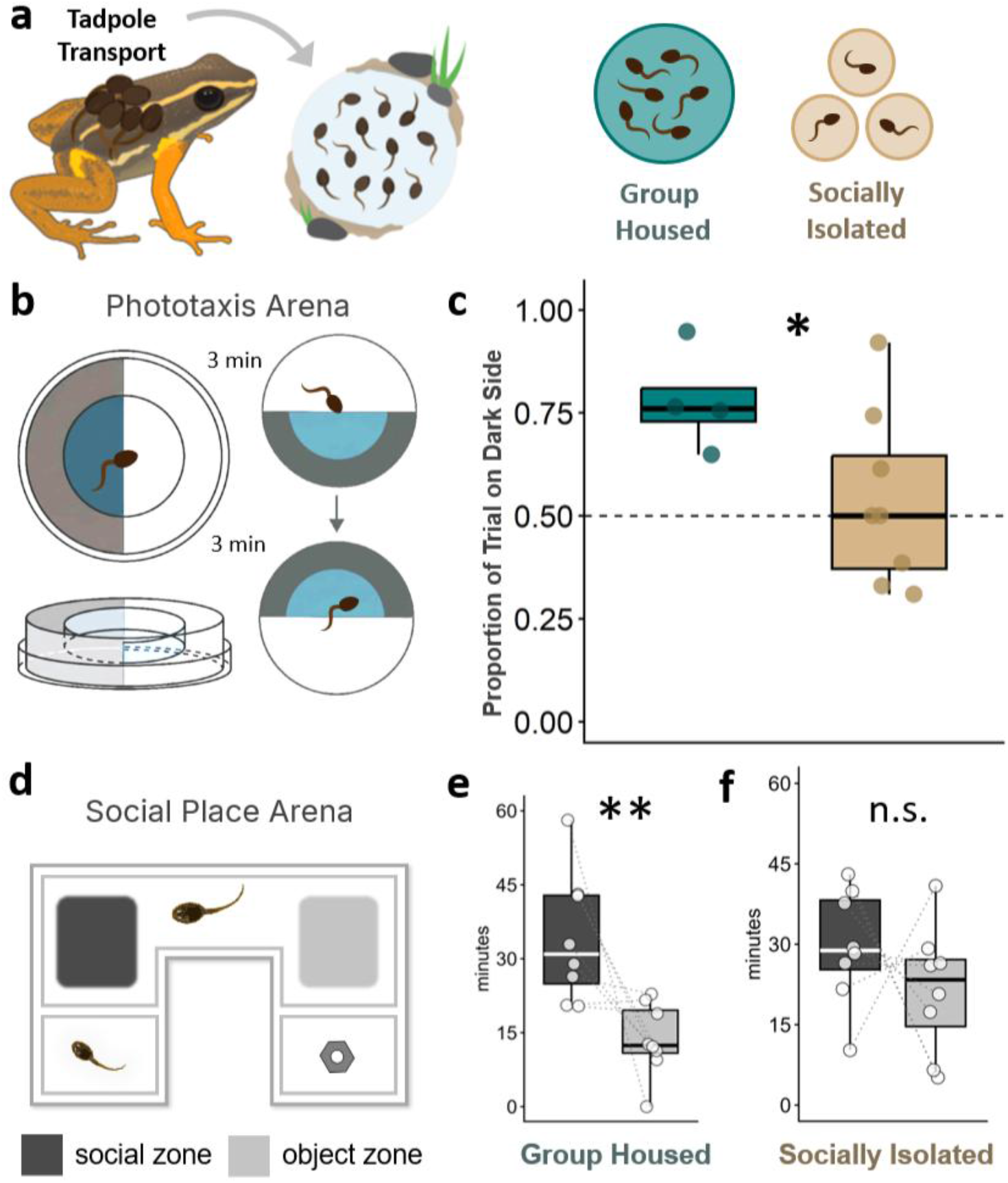
Group-housed *Allobates femoralis* tadpoles show dark side and social place preference. (a) In nature and in the lab, clutches of 12-16 tadpoles are transported by a frog to water pools. Clutches were experimentally split into either group housing (teal) or social isolation (tan) in cups. (b) Phototaxis assay: tadpoles were recorded for 3 minutes, the arena rotated 180°, and behavior recorded for a further 3 minutes. (c) Proportion of trial spent on the dark side. Group-housed tadpoles spent more time on the dark side than isolated tadpoles (Welch’s t-test, t9.5 = 2.48, p = 0.034). Points are group-housed cup means (n = 4 cups, 8 tadpoles) and individual isolated tadpoles (n = 8). Boxes show median and interquartile range; the dashed line marks chance (0.5). (d) Social place preference assay: a focal tadpole moved freely between a social zone (dark gray; near conspecific stimulus) and an object zone (light gray; near object control) separated by a neutral zone; stimulus chambers separated from arena with perforated observation barriers. (e-f) Time in each zone during the trial with perforated barriers. (e) Group-housed tadpoles spent more time in the social than the object zone (χ²1 = 11.62, padj = 0.005), whereas (f) isolated tadpoles showed no difference (χ²1 = 2.16, padj = 0.568). Points are per-tadpole; dotted lines join each animal’s social and object values (n = 8 clutches, one group-housed (GH) and one socially isolated (SI) tadpole per clutch, per assay). Boxes show median and interquartile range. (* p < 0.05, ** padj < 0.01; n.s., not significant)

**Group-housed, but not isolated, tadpoles spend more time near social stimuli** To test whether social housing affects social place preference in *A. femoralis* tadpoles, and which sensory modalities are sufficient to elicit it, we performed social place preference assays with transparent, perforated barriers, which allowed water exchange between stimulus and focal chambers, or transparent, solid barriers with visual-only access (Fig. 1d). When water exchange through the barrier was permitted, group-housed tadpoles spent more time in the social zone than in the object zone (Fig. 1e; χ²_1_ = 11.62, p_adj_ = 0.005) while isolated tadpoles showed no difference (Fig. 1f; χ²_1_ = 2.16, p_adj_ = 0.568). In visual-only trials, neither group-housed (χ²_1_ = 0.31, p_adj_ = 0.771) nor isolated (χ²_1_ = 0.46, p_adj_ = 0.771) tadpoles showed differential zone occupancy (Fig. S2). Since social rearing could also affect growth, we compared body size between isolated and group-housed tadpoles. Isolated tadpoles were larger than their group-housed clutchmates at sampling, despite stage matching and *ad libitum* feeding, exceeding them in mean per-clutch body length (paired *t-*test, t_5_ = 2.65, p = 0.046), width (t_5_ = 3.02, p = 0.029), and weight (t_5_ = 2.73, p = 0.041; n = 6 clutches; Fig. S3).

## Social rearing conditions alter miRNA expression

Since social isolation produced a change in visually guided behavior, we next tested the hypothesis that social rearing conditions would alter expression levels of microRNAs across the eye-brain axis. To do this, we compared miRNA expression profiles of group-housed and socially isolated *A. femoralis* tadpole eyes and midbrains. We chose the midbrain because this includes the optic tectum, where retinal projections from the eyes connect to the brain. Midbrain and eye libraries separated by tissue in unsupervised clustering, with distinct blocks of miRNAs distinguishing the two (Fig. S4). Within each tissue type, only one miRNA passed both significance and fold-change thresholds in a differential expression analysis between housing treatments (Fig. 2), the same miRNA in both tissues: miR-15-P1c_3p* was significantly upregulated in midbrains (log_2_FC = 0.56, p_adj_ < 0.001) and eyes (log_2_FC = 0.84, p_adj_ < 0.001).

**Figure 2.**
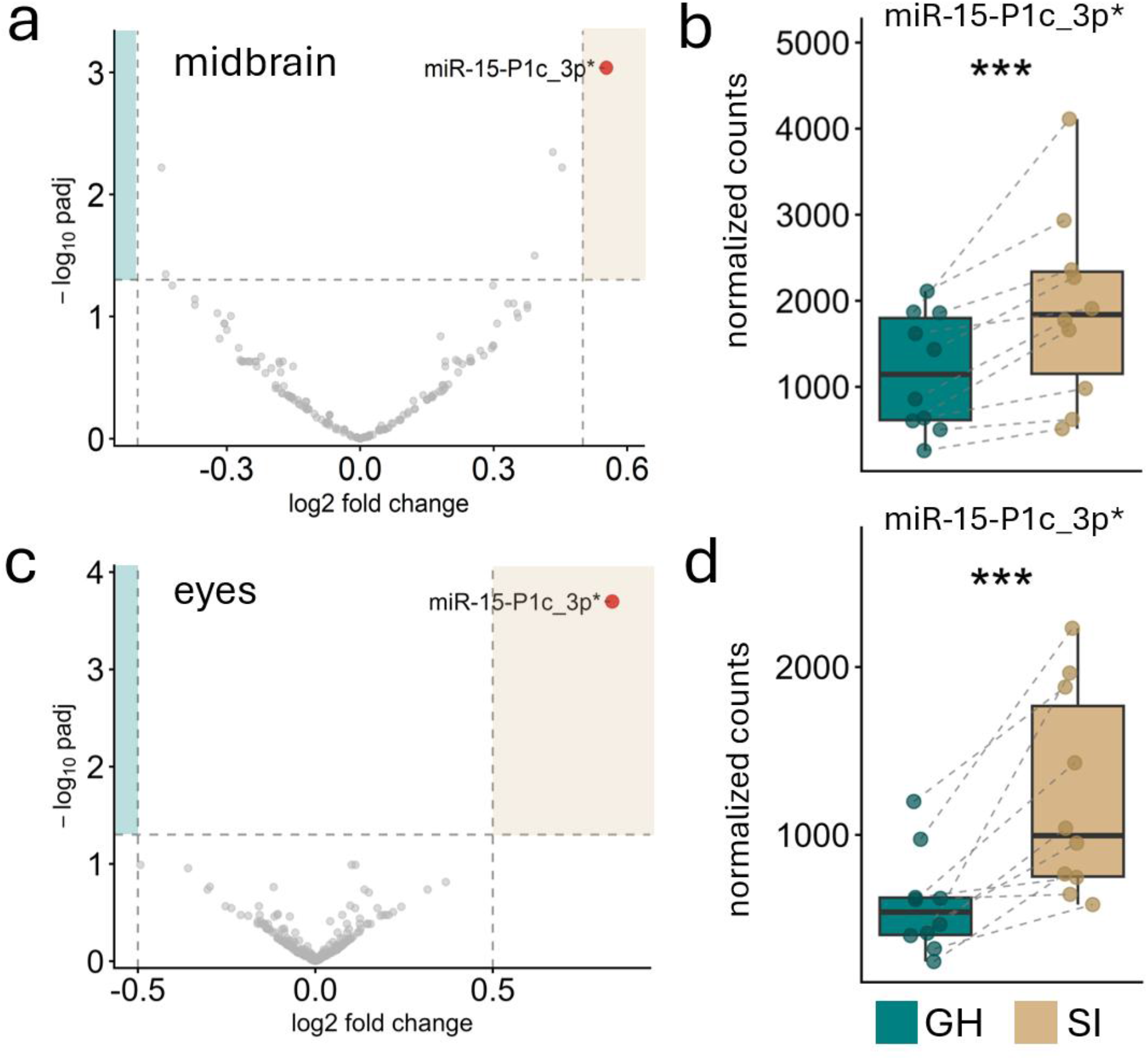
miR-15-P1c_3p* is upregulated in socially isolated midbrains and eyes. This was the only miRNA to meet significance thresholds in either (a,b) midbrain or (c,d) eyes. Left: volcano plots of miRNA differential expression (socially isolated vs. group-housed; positive log2FC indicates higher expression in isolated tadpoles) for midbrains (a) and eyes (c); dashed lines indicate significance thresholds (padj < 0.05, |log2FC| > 0.5), and the single significant miRNA is highlighted in red. Right: normalized counts of miR-15-P1c_3p* in midbrain (b) and eyes (d). Teal = group-housed (GH); tan = socially isolated (SI). DESeq2 within-tissue SI vs. GH. n = 10 clutch pairs. *** padj < 0.001.

Regulation of miRNA arm selection could yield context-dependent, alternative regulation of transcriptomes across the eye-brain axis. We next asked whether arm usage of individual miRNAs is sensitive to social rearing environments. For each tissue, we modeled the log_2_ 3p/5p arm ratio against rearing condition for every miRNA hairpin with both arms detected. Two of the 96 midbrain hairpins analyzed showed significant housing-dependent arm usage: miR-124 (Fig. 3) and miR-10-P3b (Fig. S5). No hairpin showed significant housing-dependent arm dosage differences in eyes. For miR-124 (Fig. 3a-b), the 3p arm was expressed higher than the 5p arm in both conditions, but its dominance was significantly reduced in isolated midbrains (linear model: β_SI_ = −0.73, p_adj_ = 0.035; Hedges’ g = −1.11, 95% CI [−2.16, −0.39]; Cliff’s δ = −0.66, [−1.00, −0.22]; clutch-paired Wilcoxon signed-rank, n = 10 clutch pairs, p = 0.002; Fig. 3c). miR-10-P3b shifted significantly in the opposite direction (higher 3p, β_SI_ = 0.96, p_adj_ = 0.016) but maintained a strongly 5p-dominated ratio (Fig. S5).

**Figure 3.**
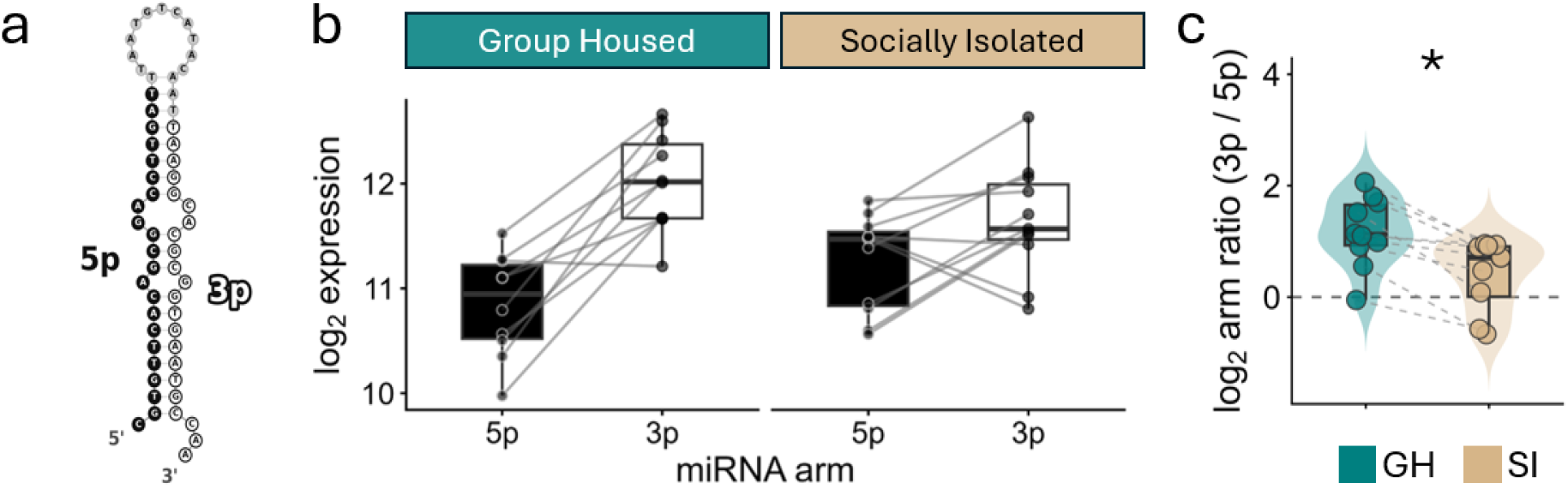
miR-124 arm dosage shifts in socially isolated midbrains. (a) Predicted secondary structure of the miR-124-P1-v1 precursor hairpin (RNAfold) (b) Paired log2 expression of miR-124 5p and 3p arms in group-housed (GH, left) and socially isolated (SI, right) tadpole midbrains; gray lines connect arms within each library. The 3p arm was expressed higher than the 5p arm in both conditions, but the 3p-5p separation narrows in isolated midbrains. (c) Per-library log2 arm ratio (3p/5p). Positive values indicate higher 3p expression, and a value of 0 represents equal arm dosage; dashed gray lines connect samples paired by clutch (Hedges’ g = −1.11, 95% CI [−2.16, −0.39]; Cliff’s δ = −0.66, [−1.00, −0.22]; clutch-paired Wilcoxon signed-rank, n = 10, p = 0.002) * padj < 0.05.

### Parathyroid hormone 2 is downregulated in socially isolated midbrains

As miRNAs can regulate the degradation of mRNA, we tested the hypothesis that miR-15-P1c_3p* would be associated with reduced expression of genes in the midbrain of socially isolated tadpoles. Only a single transcript was differentially expressed in the midbrain (Fig. 4a), while eyes showed no differential expression across housing conditions (Fig. S6). In the midbrain, *pth2* (parathyroid hormone 2) was downregulated in isolated tadpoles, relative to group-housed tadpoles (log_2_FC = −4.83, p_adj_ < 0.001) (Fig. 4b).

**Figure 4.**
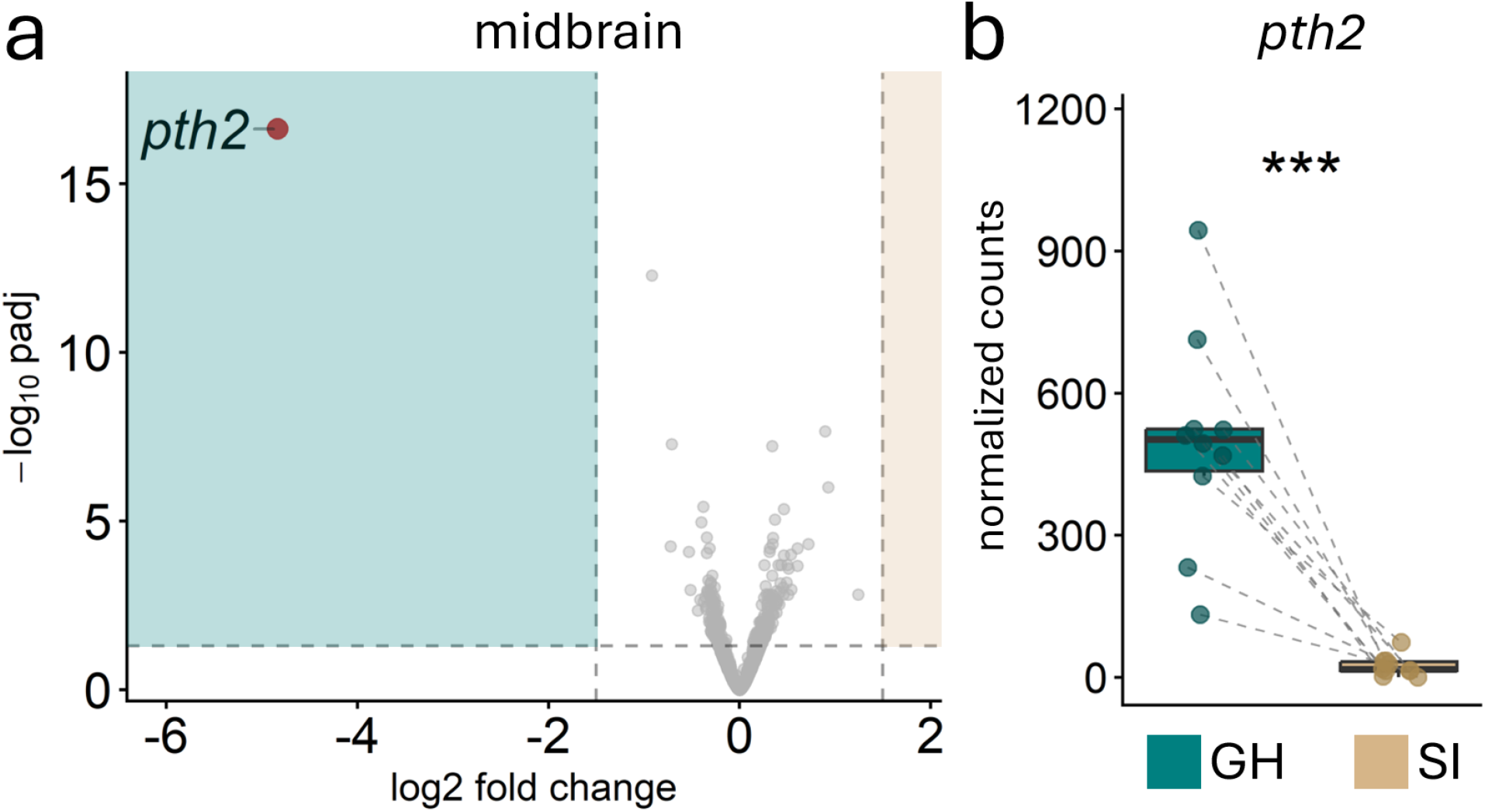
***pth2* is downregulated in socially isolated midbrains.** (a) Volcano plot of mRNA differential expression between group-housed (GH) and socially isolated (SI) midbrains. Significant hits (padj < 0.05, |log2FC| > 1.5) are red; *pth2* is the sole transcript passing both thresholds. Dashed lines mark significance and fold-change cutoffs. Shaded panels indicate direction of enrichment (teal, GH; tan, SI). (b) *pth2* normalized counts by housing condition. Points are paired within clutch, n = 10 pairs. *** padj < 0.001.

As isolated tadpoles had increased miR-15-P1c_3p* and decreased *pth2* expression in midbrains, we next attempted to detect miR-15-P1c_3p* binding sites in the *pth2* 3’UTR. However, the *A. femoralis* genome had no annotated *pth2* 3’UTR. We recovered the 3’UTR from our own RNA-seq coverage, which placed the 3’ end 81 nt past the stop codon. This predicted *A. femoralis* 3’UTR contained no binding site for miR-15-P1c_3p*, nor either arm of miR-124.

## Discussion

Regulatory programs respond to developmental conditions to enable context-dependent reshaping of the transcriptome. In the nervous system, microRNAs can regulate gene expression in response to social shifts and thereby alter developmental trajectories and behavioral plasticity. The brilliant-thighed poison frog, *Allobates femoralis*, exhibits behavioral plasticity across life stages, including flexible parental care in adults (Ringler et al., 2015) and social rearing as tadpoles. Our comparisons of brain and eye miRNA profiles and transcriptomes between group-housed and socially isolated tadpoles revealed the impact of early-life isolation on behavior and miRNA expression in key parts of the developing *A. femoralis* visual system.

## Early-life isolation alters dark side preference behavior

Light/dark phototaxis assays are used as a proxy for exploratory and anxiety-like behaviors in many species (Bourin and Hascoët, 2003; Maximino et al., 2010). We found that group-housed *A. femoralis* tadpoles preferred dark environments while isolated tadpoles showed no light/dark preference. Phototactic behaviors vary across amphibian species. *Xenopus* tadpoles exhibit a light-side preference, and when the optic nerve is severed, this preference is abolished (Viczian and Zuber, 2014). Tadpoles of the mimic poison frog, *Ranitomeya imitator* (Schulte, 1986), exhibit dark side preference, similar to *A. femoralis* tadpoles, and this preference is maintained even when blinded (Butler et al., 2024). Thus, dark side preference appears in at least two poison frog species, but its sensory basis and ecological significance may differ from the light preference observed in *Xenopus* tadpoles.

Group rearing may alter visual experience directly. Visual cues from conspecifics increase swimming activity in *A. femoralis* tadpoles (Szabo et al., 2021), suggesting that social isolation could impact general motor behaviors through deprivation of visual cues provided by conspecifics. This could also impact development of visual circuits linked to predator avoidance, as *A. femoralis* tadpoles combine visual and chemical cues to avoid dragonfly larvae, a natural predator (Szabo et al., 2021). Relatedly, in a shelter-emergence assay, tadpoles originating from natural pools containing dragonfly larvae show longer initial emergence latency and greater unpredictability in emergence timing, compared to predator-naïve tadpoles _(McPherson_ et al., 2026). Thus, early developmental environment impacts tadpole behavior, with consequences ranging from gross conspecific responses to extended sheltering post-predator exposure. That isolated *A. femoralis* tadpoles do not present dark side preference here is suggestive of a role for conspecific cues in the development of broad responses to light/dark contrast, with possible impacts on risk-related behavior.

## Social place preference depends on water exchange

Social place preference assays measure an animal’s sensitivity to social stimuli. Developmental social isolation alters social preference across vertebrates, including zebrafish (Shams et al., 2018) and mice (Yamamuro et al., 2020). We found that group-housed *A. femoralis* tadpoles preferred the social zone when stimulus chambers were separated by a perforated barrier, whereas isolated tadpoles showed no preference. Neither group showed any preference when only provided visual access to stimuli. Since tadpole visual systems are immature at this developmental stage, animals may rely more on non-visual cues to detect conspecifics (Butler et al., 2024). The transparent, perforated barrier permits both chemical and water movement cues to pass to the focal chamber. Social preference in group-housed tadpoles could therefore reflect chemosensory or mechanosensory (e.g. lateral-line mediated) detection, or some combination of the two, alongside vision. A reliance on waterborne cues would align with the ecology of this species, as adult *A. femoralis* use waterborne odor cues to locate suitable pools for tadpole deposition (Serrano-Rojas and Pašukonis, 2021).

This contrasts with prior work with *A. femoralis* tadpoles, where visual cues from conspecifics were sufficient to alter tadpole swimming activity (Szabo et al., 2021). That work measured general activity rather than social proximity, whereas we compared spatial preference between two groups reared in divergent social settings. Notably, that study also found that visual and chemical cues combined were sufficient to elicit avoidance of dragonfly larval predators, indicating that sensory modality requirements for *A. femoralis* tadpoles can be behavior-specific. Future work could investigate whether group-housed and isolated *A. femoralis* tadpoles have different gross swimming activity in social stimulus assays.

## Social rearing environment alters expression patterns of microRNAs

Social rearing conditions altered expression patterns of microRNAs, including abundance and the relative dosage of the dominant arm. Isolated tadpoles exhibited higher levels of miR-15-P1c_3p* in both eyes and midbrain. The miR-15 family is well characterized in mammals, where the 5p arm has been linked to cancers, neurodegenerative disease, and neuroinflammation (Finnerty et al., 2010). Notably, another paralog (miR-15-P1a) is documented to be involved in neuroplasticity, modulating dendritogenesis by regulating *bdnf* expression in developing neurons (Gao et al., 2015). However, that work does not extend to the 3p arm.

The miR-15-P1c locus is conserved across vertebrates (Clarke et al., 2025) and its 5p arm carries the deeply conserved miR-15 family seed (AGCAGCA) in both frogs and mammals. While the 3p* arm only differs between *X. tropicalis* and *A. femoralis* by a single position (9, U → C), with an identical seed, it shares only 50% sequence identity with the human ortholog (differing at 3/7 seed positions), so predicted mammalian targets for this arm do not transfer. As the response is shared across eyes and midbrain, future work could investigate its spatial expression to test whether it regulates retinotectal development and how any regulation may influence sensorimotor behaviors.

Since alternate arms of the same miRNA hairpin carry different seed sequences (and therefore different target sets), a shift in relative arm output may redistribute regulation across two transcript sets, rather than simply raising or lowering one set in isolation (Pinhal et al., 2025). We found two miRNAs that showed housing-dependent arm ratio shifts: miR-10-P3b and miR-124. For miR-10-P3b, isolation resulted in a significant shift, but the 5p arm still accounted for the majority of reads in both conditions. For miR-124, isolation shifted arm expression toward comparable representation between the two arms, indicating a change in arm dosing. miR-124 is among the most abundant miRNAs in the mammalian brain (Lagos-Quintana et al., 2002), contributing to various neural processes including neurite outgrowth, differentiation, neurotransmission, and synaptic morphology, among others (Rajasethupathy et al., 2009; Sun et al., 2015). In the developing optic tectum of *Xenopus* tadpoles, miR-124 gates the timing of growth cone sensitivity to guidance cues during retinotectal pathfinding (Baudet et al., 2012).

Previously, miR-124 was linked to depression-like behavior (Roy et al., 2017), social dysregulation (Bahi, 2017), and stress responses to social isolation (Pan-Vazquez et al., 2015), but in each case, the reported changes are in the abundance of just the dominant arm. To our knowledge, this is the first report of a socially dependent change in miR-124 arm dosing. The shift was restricted to the midbrain (unlike miR-15-P1c_3p*, which responded in both tissues), placing it in the tissue where retinotectal circuits are assembled and where the transcriptome also responded to isolation.

Resolving which transcripts are affected would require annotated 3’UTRs, which the current *A. femoralis* annotation does not provide. Future work with an improved genome could reveal if networks of genes were impacted by this regulatory shift.

## *pth2* downregulation recapitulates a conserved signature of social isolation

The only transcript differentially expressed between social rearing groups was in the midbrain: parathyroid hormone 2 (*pth2*), also known as tuberoinfundibular peptide of 39 residues, or TIP39. *pth2* was downregulated in isolated *A. femoralis* tadpoles, similar to studies in zebrafish (Anneser et al., 2020). Synthesis of *pth2* is restricted to the posterior thalamus and lateral pons in rodents (Dobolyi et al., 2012), and the thalamus of zebrafish (Bhattacharya et al., 2011). These sources lie immediately anterior and posterior to our midbrain dissection, so we cannot exclude a contribution from adjacent tissue in our libraries.

Zebrafish with a *pth2* knockout show intensified startle response, less cohesive shoaling, and attenuated social place preference (Anneser et al., 2022). Additionally, *pth2* is linked to mechanosensation, where ablation of the lateral line in isolated animals prevents the *pth2* recovery that normally accompanies social re-exposure, and artificially generated water movement is sufficient to restore *pth2* levels in the absence of social re-exposure (Anneser et al., 2020). These zebrafish findings align with our behavioral results, where social place preference in *A. femoralis* tadpoles emerged when water exchange between the social chamber and focal arena was permitted, implicating chemosensory and/or mechanosensory cues. This peptide is unlikely to be involved in phototaxis differences here, as *A. femoralis* tadpole light/dark preferences were assayed individually and in the absence of conspecific cues. Future work could involve testing whether *pth2* signaling mediates social place preference in *A. femoralis*, using pharmacology or lateral line ablation.

Though we did not detect binding sites for miR-15-P1c_3p* in our manual curation of the *pth2* 3’UTR, it is possible that other transcripts with sub-threshold expression shifts were impacted by the miR-15-P1c_3p* upregulation in isolated samples. A major limitation of this study is that mRNA targets of the microRNAs associated with social rearing conditions in *A. femoralis* are unknown, as the genomic resources for this species are currently underdeveloped. However, with an improved genome, this analysis will be possible in future studies. Isolated tadpoles also grew larger than their age- and stage-matched clutchmates, despite *ad libitum* feeding. Growth and the expression differences we report are both downstream of the experimental social manipulation, and our design cannot resolve whether isolation acts on the eye-brain axis directly, or indirectly through the growth differences it produces.

## Conclusion

This work leveraged the natural plasticity of a poison frog tadpole brain and behavior to examine regulatory expression correlates of social environment shifts. Though it is broadly accepted that miRNAs play outsized roles in the developing brain, knowledge of specific brain miRNA correlates of social shifts is just beginning to surface. We showed that social environment shaped miRNA expression and arm dosage, in tandem with behavioral and transcriptional shifts. Future studies should include functional manipulations to test the role of these miRNAs in developmental processes.

Additionally, comparative experiments would be informative, as rearing conditions also vary naturally across the dendrobatid tree, from solitary deposition to communal pools. Comparing the magnitude of isolation-induced miRNA responses across these closely related (but behaviorally divergent) species could reveal whether regulatory social sensitivity tracks the evolution of their social behavior.

## Acknowledgements

We thank members of the O’Connell lab, especially Madison Lacey and Dr. Mesi Fischer, for their assistance with animal care. We also thank Dr. Roberto Márquez and Dr. Peter Sarnow for discussions about data analysis, and Dr. Daniel Shaykevich and Dr. Mesi Fischer for comments.

## Data and resource availability

Raw sequencing reads will be deposited in the NCBI Sequence Read Archive and accession numbers provided upon acceptance.

## Funding

This work was supported by National Institutes of Health grants (R01HD110514 and R56MH133094) to LAO.

## Competing interests

The authors declare no competing or financial interests.

## Ethical statement – IACUC

All animal procedures were approved by the Stanford Administrative Panel on Laboratory Animal Care (protocol no. 33097).

## Author contributions

Conceptualization: NMK, JMB, LAO Methodology: NMK, JMB, LAO Validation: NMK

Formal Analysis: NMK, MMM Investigation: NMK, MMM, KN, MGP, JMB Resources: LAO

Data Curation: NMK

Writing – original draft preparation: NMK

Writing – review and editing: NMK, LAO, MMM, KN, MGP, JMB Visualization: NMK, LAO

Project Administration: NMK, LAO Supervision: LAO

Funding Acquisition: LAO

## Supplementary Figures

**Fig. S1:**
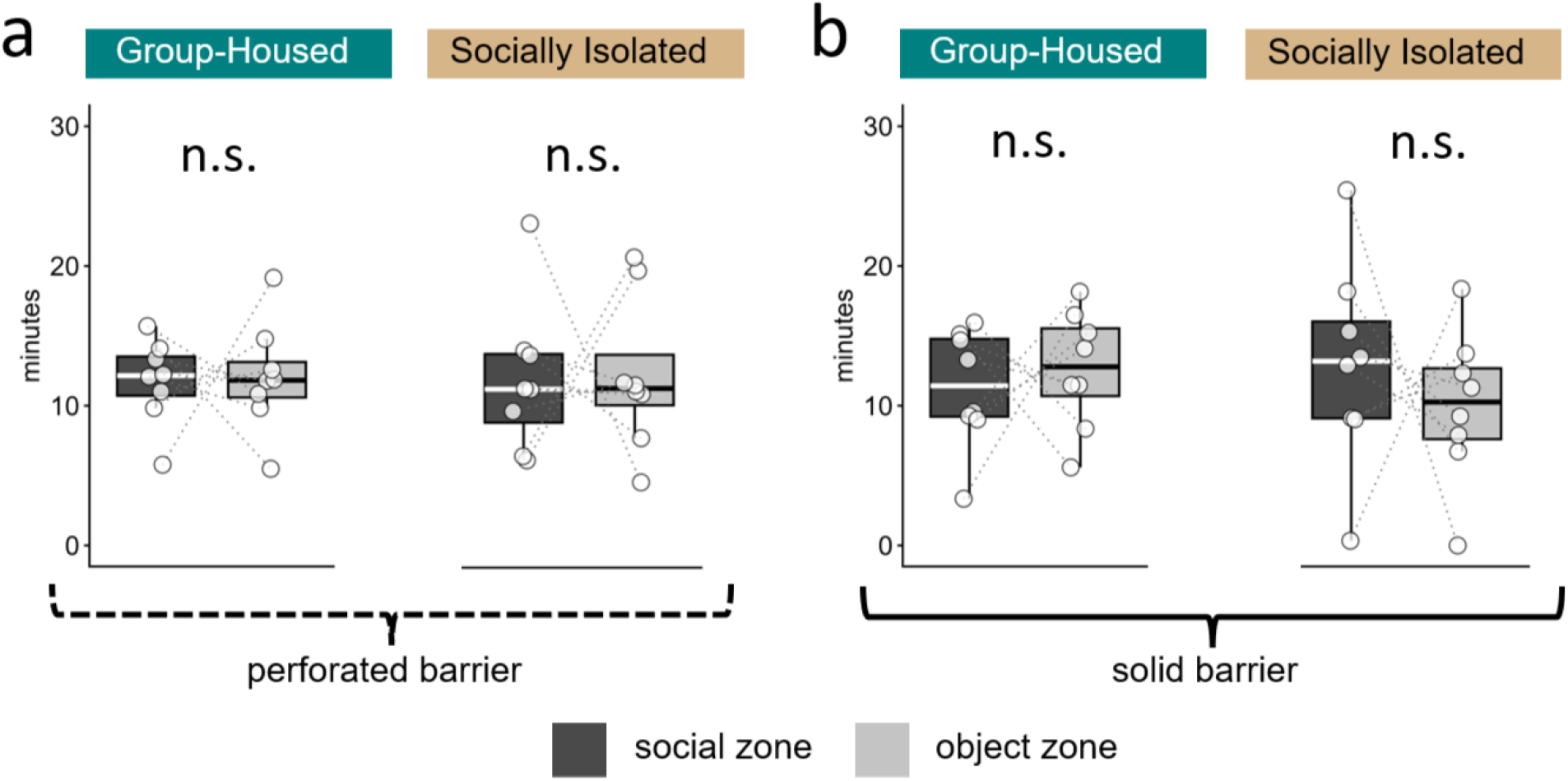
Tadpoles show no zone preference during the baseline acclimation period. Time in the social (dark gray) and object (light gray) zones during the 30-minute acclimation phase before stimuli introduced. Panels show barrier types separating focal tadpoles from stimuli (a, perforated, permitting chemical/visual/water exchange; b, solid, visual-only access), split by social housing condition (group-housed, socially isolated). Dotted lines connect zone values from the same tadpole. Boxes show median and interquartile range. No group differed between zones. (a) perforated barrier (chem/visual/water access): group-housed (χ²1 = 0.03, padj = 0.907), socially isolated (χ²1 = 0.01, padj = 0.907). (b) solid barrier (visual-only access): group-housed (χ²1 = 0.44, padj = 0.771), socially isolated (χ²1 = 0.97, padj = 0.771). Linear mixed model with a random intercept for tadpole; p-values Benjamini-Hochberg corrected across all barrier x rearing x phase comparisons (n = 8 clutches, one group-housed and one isolated tadpole per clutch, per barrier type). n.s., not significant.

**Fig. S2:**
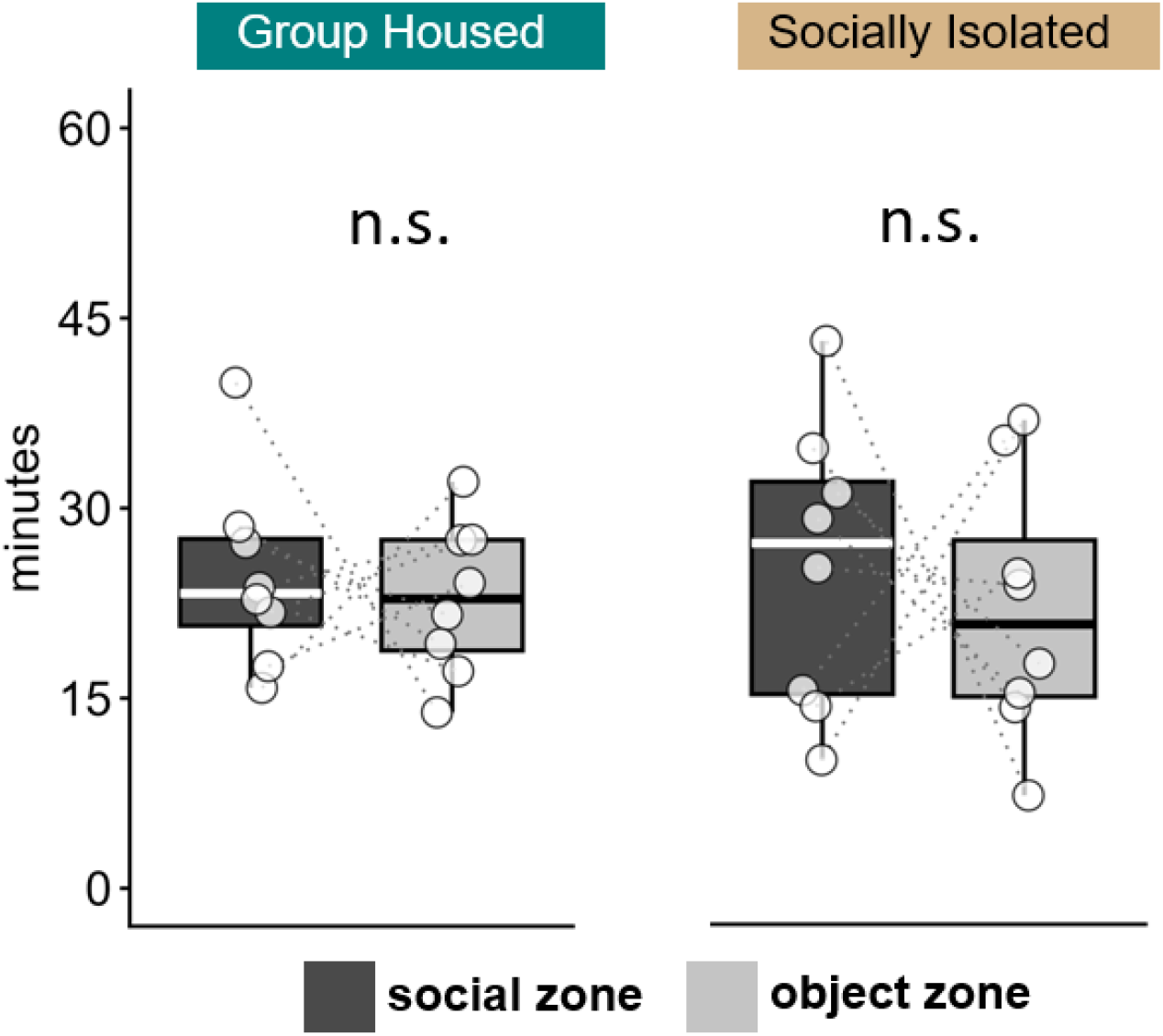
Time in social place preference zones with visual-only access. Social place preference trials with solid, transparent barriers permitting visual access without water exchange. Time (minutes) in social (dark gray) and object (light gray) zones for group-housed (left) and socially isolated (right) tadpoles. Dotted lines connect occupancy values from the same tadpole. Neither group-housed (χ²1 = 0.31, padj = 0.771) nor isolated tadpoles (χ²1 = 0.46, padj = 0.771) differed between zones. Separate animals from Fig. 1: one GH and one SI per clutch (n = 8 clutches). Boxes show median and interquartile range.

**Fig. S3:**
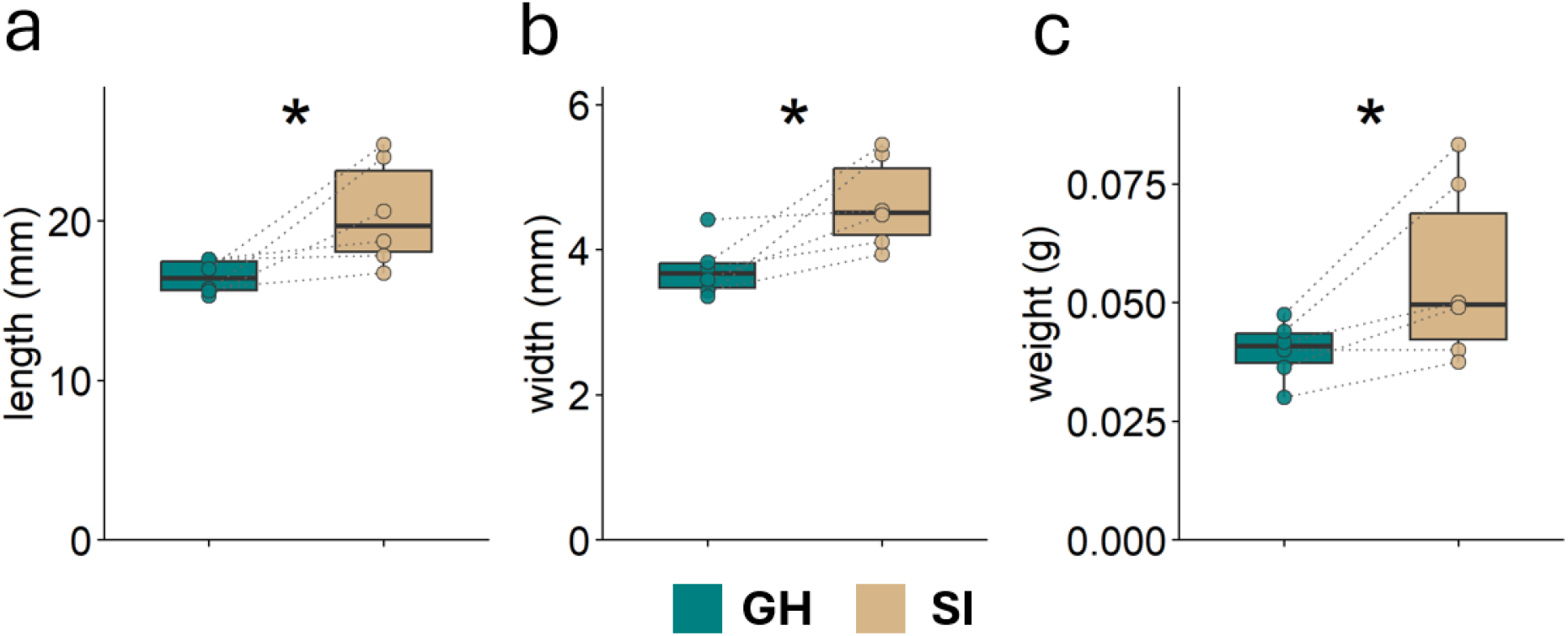
Socially isolated tadpoles are larger than stage-matched group-housed clutchmates. Size comparison of group-housed (GH, teal) and socially isolated (SI, tan) *A. femoralis* tadpoles. (a) length (t5 = 2.65, p = 0.046), (b) width (t5 = 3.02, p = 0.029), and (c) weight (t5 = 2.73, p = 0.041). Points are clutch/condition means (n = 6 clutches; dotted lines connect paired clutches, * p < 0.05).

**Fig S4:**
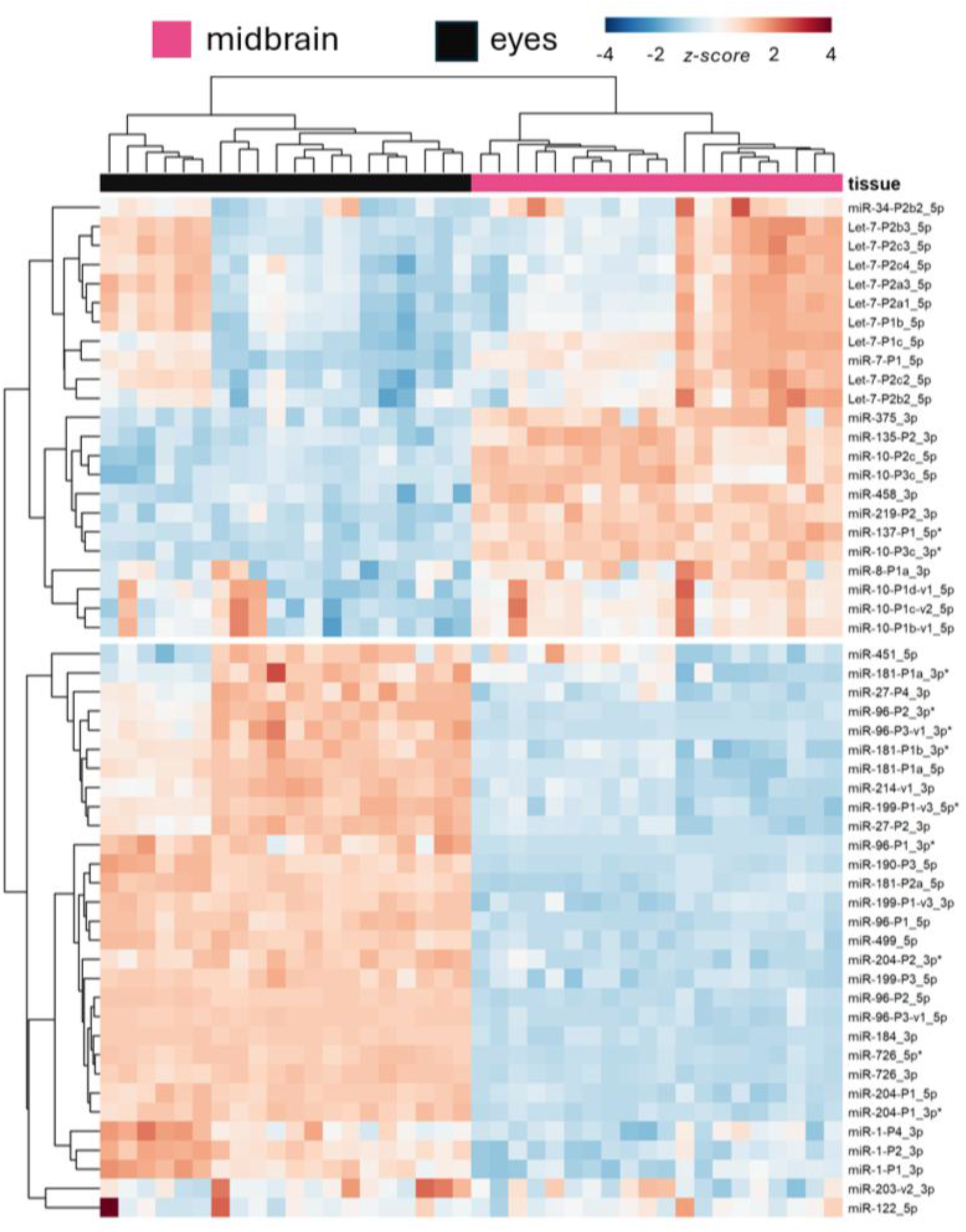
Unsupervised clustering separates libraries by tissue. Heatmap of the top 20% most variable miRNAs (53 miRNAs) across midbrain (n = 20, pink) and eye (n = 20, black) libraries. Values are variance stabilized counts (DESeq2) centered and scaled to unit variance by row (z-score; color bar). miRNA names follow MirGeneDB naming convention; * denotes the non-canonical arm.

**Fig. S5:**
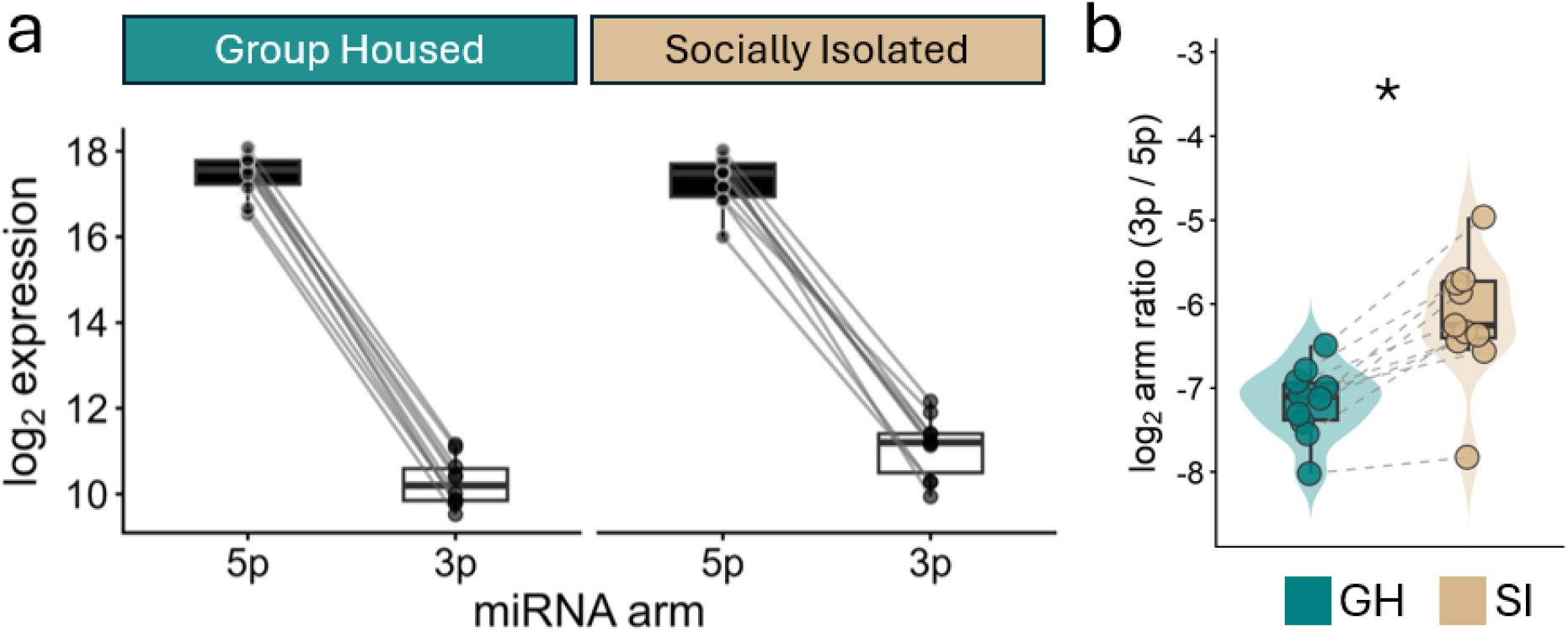
**miR-10-P3b arm dosage shifts in socially isolated midbrains**. (a) Paired log2 expression of miR-10-P3b 5p and 3p arms in group-housed (teal, left) and socially isolated (right, tan) tadpole midbrains. Gray lines connect arms within each midbrain library. The 5p arm was expressed far higher than the 3p arm in both conditions, and this separation slightly narrows in isolated midbrains. (b) Per-library log2 arm ratio. Negative values indicate higher 5p expression. Unlike miR-124 (Fig. 3) the ratio remains highly 5p dominant across social conditions (linear model, βSI = 0.96, padj = 0.016; Hedges’ g = 1.52, 95% CI [0.63, 3.26]; Cliff’s δ = 0.80, [0.38, 1.00]; clutch-paired Wilcoxon signed-rank, n = 10, p = 0.002). * padj < 0.05.

**Fig. S6:**
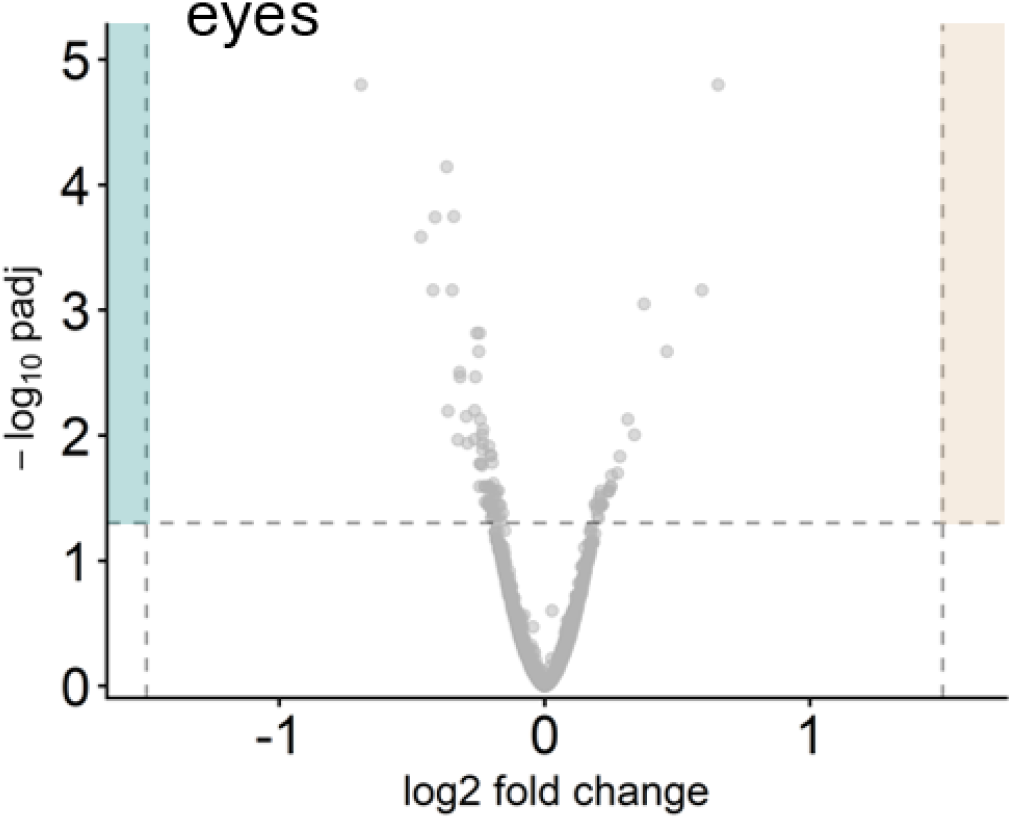
**No transcript is differentially expressed between housing conditions in eyes**. Volcano plot of mRNA differential expression between group-housed (GH) and socially isolated (SI) eyes. Dashed lines mark significance and fold-change cutoffs (padj < 0.05, |log2FC| > 1.5).

## Notes

### Competing Interest Statement

The authors have declared no competing interest.

